# A metabolite extracted from *E. coli* suppresses tau aggregation

**DOI:** 10.1101/2025.10.30.684975

**Authors:** Mahalashmi Srinivasan, Akhil Patel, Tark Patel, Jack Moore, Erik Gomez-Cardona, Allan Yarahmady, Brian D. Sykes, Olivier Julien, Sue-Ann Mok

**Author notes:** equal contribution (Akhil Patel, Tark Patel).

## Abstract

Tau aggregation is a key pathological feature of neurodegenerative diseases termed tauopathies. Identifying the various cellular factors that function to prevent tau aggregation in cells can generate key insights into how to mitigate diseases associated with protein misfolding. During an investigation into developing purification methods for the protein tau, we observed that isolates of *E. coli* lysate prevented human tau aggregation *in vitro*. Fractionation of the lysate was used to further isolate a small molecular weight (MW) inhibitory fraction containing multiple components, as determined by mass spectrometry and NMR. A putative inhibitory component, methylphosphonic acid (MePn), decreased tau amyloid formation when supplemented to *in vitro* aggregation assays. MePn also blocked the aggregation of expressed tau in live *E. coli* when supplemented to the culture media. Our findings can be directly applied to optimizing purification of recombinant tau protein and more broadly, highlight the potential of cellular metabolites to directly modulate tau amyloid formation.

## Introduction

Accumulation of amyloid aggregates composed of the microtubule associated protein tau (commonly referred to as tau) is a pathological hallmark of neurodegenerative diseases classified as tauopathies (1, 2). An example of tauopathy is Alzheimer’s Disease which is characterized by two distinct amyloid protein deposits: tau and amyloid beta (3, 4). Amyloid beta aggregates form earlier in Alzheimer’s disease progression; however, tau aggregate accumulation is more strongly correlated with clinical symptoms of cognitive decline (5). In another tauopathy, Frontotemporal Lobar Degeneration, tau can be associated with mutations in the microtubule associated protein tau (MAPT) gene that encodes tau (6). These and other lines of evidence support the current hypothesis that tau dysfunction related to its aggregation promotes neurodegenerative disease.

Tau is an intrinsically disordered protein (7–10) primarily expressed in neurons with an axonal and soma distribution (11). Tau has well-characterized interactions with tubulins and their assembled microtubule forms (12, 13). Many other tau interaction partners have also been detected that could potentially modulate pathology development (14–18). Soluble tau has ascribed functions in regulating multiple cellular processes such as microtubule dynamics and stability, axonal transport, mitochondrial health and myelination (19). In contrast, tau molecules can self-assemble into pathogenic higher-order aggregated species in the form of oligomers and amyloidogenic fibrils (20–23). Aggregated tau species can transmit from cell-to-cell and template new rounds of tau aggregation thereby promoting the amplification and spreading of pathology and cellular dysfunction (24).

The identification of cellular factors that regulate the conversion of tau from its soluble to aggregated forms is an intense area of investigation. For example, changes in protein-protein interaction partners such as microtubules (25) or molecular chaperones (26) have been proposed to increase the probability of tau to aggregate. Alterations in the biochemical nature of tau via post-translational modifications have also been observed to regulate tau aggregation. In general, hyperphosphorylation of tau promotes its aggregation (27, 28). In contrast, tau glycosylation inhibits its aggregation (29). Tau aggregation has also been shown to be modulated by biological molecules such as RNA (30), polyphosphates (31), and catecholamines (norepinephrine, epinephrine) (32, 33). It is probable that other not yet identified cellular components also contribute to maintaining the solubility of tau in healthy cells.

We found evidence of tau aggregation inhibitor activity in *E. coli* cell lysates and we set out to isolate the inhibitory molecule(s). Our isolation procedures led us to identify the metabolite methylphosphonic acid as a suppressor of tau aggregation in *in vitro* and *in cellulo* models.

## Materials and Methods

### Inhibitor fractionation from *E. coli*

BL21(DE3) RP *E. coli* cells (NEB) transformed with pUC19 or pET28 0N4R human tau (34) were inoculated into a starter culture of Terrific Broth (TB) containing ampicillin (100 µg/mL) or kanamycin (50 µg/mL), respectively. The culture was incubated overnight (37°C, 250 rpm) then transferred at a 1:50 dilution to 1 L of fresh TB containing appropriate antibiotics. Cultures were incubated with shaking (37°C, 250 rpm) until OD600 = 0.6-0.7. For cells transformed with pET28 0N4R tau, 500 µM isopropyl-beta-D-thiogalactopyranoside (IPTG) was added to induce protein expression and incubated for an additional 3 hrs. Cells were collected by centrifugation (5000 x g, 15 min) and resuspended in resuspension buffer [D-PBS, pH 7.4, [1X] Complete protease inhibitor cocktail (Roche), 1 mM phenylmethylsulfonylfluoride (PMSF), 2 mM dithiothreitol (DTT)]. Bacteria were lysed with lysozyme (100 mM Tris pH 8.0, 50% glycerol) at a final concentration of 0.3 mg/mL over 4 freeze-thaw cycles. The lysate was treated with 11.6 units/mL benzonase (sigma) for 20 min at 4°C, boiled for 20 min to denature proteins, then centrifuged at 19,500 x g for 45 min. The supernatant was passed through a 0.22 µM MCE filter (Millipore) then separated with a HiLoad 16/600 Superdex 75 prep column (Cytiva) equilibrated with D-PBS and 2 mM DTT. Eluted fractions (1 mL) were collected and stored at −80°C until further use.

### Tau protein purification

Large-scale purifications of WT 0N4R WT human tau protein and PAD12 (0N3R and 0N4R) constructs were carried out as previously described in Mok *et al*. (34). The WT 0N4R sequence was previously cloned into pET28a vector (34). The sequences for the phosphomimetic mutants, 0N4R and 0N3R PAD12 (35) were cloned into the pET-IDT vector for this study. Briefly, tau expression was induced in BL21(DE3)RP *E. coli* transformed with the desired expression constructs (TB, 50 µg/mL Kan, 500 µM IPTG) for 3 hrs at 30°C. Cells were lysed with a high pressure homogenizer in resuspension buffer (20 mM MES, pH 6.8, 1 mM EGTA, 0.2 mM MgCl_2_, 1 mM PMSF, Complete protease inhibitor cocktail, 5 mM DTT), boiled for 20 min after adding NaCl to a final concentration of 0.5 M, then centrifuged (19,500 x g, 45 min). The supernatant was dialyzed overnight in Cation A buffer (20 mM MES pH 6.8, 50 mM NaCl, 1 mM EGTA, 1mM MgCl_2_, 100 mM PMSF, 2 mM DTT) and processed by cation exchange chromatography (elution gradient: 150-600 mM NaCl). Eluted fractions containing purified tau were identified by coomassie-stained SDS-PAGE then pooled, concentrated and exchanged into aggregation assay buffer (D-PBS, pH 7.4, 2 mM DTT) prior to storage at −80°C.

### Tau aggregation assay

0N4R WT tau (10 µM) in 0.22 µm filtered aggregation assay buffer (D-PBS, pH 7.4, 2 mM DTT) was aggregated following the addition of 88 μg/mL heparin (Santa Cruz) at 37°C with shaking as previously described (36). Similarly, 0N4R and 0N3R PAD12 tau in D-PBS (pH 7.4, 2 mM DTT) was aggregated at a final concentration of 40 µM in the absence of any inducers at 37°C with shaking. For equimolar mixtures of 0N4R and 0N3R PAD12 tau, 20 µM of each protein was present in the final reaction. When fractionated inhibitor samples were added to the aggregation assay, buffer only matched control reactions were run in parallel. Methylphosphonic acid (Sigma) was prepared as a stock solution in aggregation assay buffer which was adjusted to a final pH of 7.4 prior to use. For kinetic assays, 10 µM of the amyloid binding dye thioflavin T (ThT) was also included in the reactions (20 µL total) which were set up in 384 well, low-volume,plates. To monitor amyloid formation, ThT fluorescence was measured every 5 min in a SpectraMax M3 plate reader (Molecular Devices) with the following settings: λex=444 nm, λem= 485 nm, cut-off=475 nm, 37°C, continuous shaking between reads. For analysis of aggregation kinetics, individual aggregation curves were fitted to the Gompertz function as described previously (36)to extract the kinetic parameters lag time and amplitude. Analysis was performed on baseline subtracted curves unless otherwise noted in the text.

### Tau aggregate sedimentation assay

Aggregation reactions were carried out in 200 µL reaction volumes in low binding 1.7 mL microfuge tubes (Denville) at 37°C for 24 hrs with continuous shaking (800 rpm). Tau aggregates were isolated from completed reactions by centrifugation at 100,000 x g for 1 hr. The supernatant was removed without disturbing the pellet, and pellets were resuspended in 20 µL D-PBS before being transferred to non-binding tubes for storage at −80°C. Supernatant and pellet samples were separated on a 4-20% SDS-PAGE gel. ImageJ was used to determine the densitometry profile of each sample.

### Trypsin digestion of protease-resistant fragments for tau aggregates

Samples were processed as previously described (36). Tau aggregation reactions were incubated with mass spectrometry grade trypsin (ThermoFisher) at a final concentration of 29 µg/mL, at 37°C for 1 hr with shaking (800 rpm). The digested reaction samples were immediately stored at −80°C until further processing by capillary gel electrophoresis (ProteinSimple) using the total protein detection module. Briefly, digested aggregate reactions were mixed 5:1 with ProteinSimple 5X sample buffer then boiled for 5 min at 96°C. Samples were resolved on 2-40 kDa cartridges (ProteinSimple) following manufacturer protocols.

### HPLC separation of tau inhibitor fractions

HPLC was performed using an Agilent 1200 series HPLC. The sample was loaded onto a Luna Omega C18 HPLC (00F-4753-AN, Phenomenex) at a flow rate of 250 μL/min, a column temperature of 40°C, and a buffer composition of 2% B (A = water + 0.01% TFA, B = acetonitrile + 0.1% TFA). The composition was held at 2% B for 5 min and then increased to 82% B over 40 min while monitoring absorbance at both 210 nm and 260 nm.

### NMR acquisition and data processing

The NMR experiments were all performed on a Varian 500 MHz INOVA NMR spectrometer. The one-dimensional (1D) proton NMR spectra were acquired with 128 transients, at 25°C. The processing and visualization of the 1D spectra was done with the VnmrJ software v2.1B (Varian inc.). Each spectrum was processed with a line broadening of 1.5 Hz. The concentrations of MePn were determined by relative peak heights with respect to the blank (standard spectrum) under the same conditions.

### Mass Spectrometry of *E. coli* isolates and MePn

The direct infusion measurement was carried out on an Q Exactive Orbitrap mass spectrometer (Thermo Scientific) using the HESI source and the on-board syringe pump at a flow rate of 10 μL/min. The following parameters were used: resolution of 70,000, AGC target of 1e6, maximum inject time of 250 ms, spray voltage of 3.0 kV, capillary temperature of 275°C, S-lens RF of 60.0, sheath gas flow of 10. Mass calibration was done externally by direct infusion of a standard ion calibration solution (LTQ Velos ESI Positive Ion, Thermo Scientific).

### Thioflavin-S detection of amyloids in *E. coli*

For overnight culture preparation, 10 mL of M9 minimal medium containing 50 μg/mL of kanamycin was inoculated with a single colony of BL21 (DE3) bearing the plasmids for expression of 0N4R human tau or His-MBP-46Q htt exon 1 at 37°C. The overnight culture was diluted 1:10 in fresh M9 minimal medium (5 mL) containing kanamycin (50 μg/mL), Thioflavin-S (ThS) (25 μM), and different concentrations of MePn (16-130 mM). The cultures were grown to an OD600 between 0.6 to 0.8 and 1 mL samples were collected for uninduced controls. Protein expression was induced with 0.5 mM IPTG, at 37°C and shaking (250 rpm) for 18 hrs. For a negative control group, water was substituted for MePn addition. Amyloid formation was measured and analyzed as described in (37). Briefly, 200 μL samples were aliquoted to a 96-well plate in triplicates after normalizing for cell density (OD600). Samples were excited at 440 nm and the fluorescence emission spectra was collected from 475-600 nm (emission cut off = 475 nm).

Emission spectra values were processed by background subtracting the values from uninduced samples.

### Western Blot

Equal cell densities (based on OD600 measurements) of uninduced and induced cultures were pelleted at 3,000 x g for 10 min. Cell pellets were resuspended in 20 µL of water with 4x sample buffer, and boiled at 95°C for 10 min. The boiled samples were centrifuged at 13,000 x g for 15 min, and the supernatants were loaded onto a 4-20% Tris-glycine SDS PAGE gel. Proteins were transferred to a 0.1 µM nitrocellulose membrane (Cytiva Amersham), which was then blocked with 2.5% fish skin gelatin dissolved in 1x TBST. Blot was incubated with primary antibodies against D1M9X (Cell Signaling Technology) for 0N4R tau and 6x His Tag (Thermo Fisher) for His-MBP-mutant Htt fusion protein. HRP-conjugated secondary antibodies (Cell Signaling Technology) and chemiluminescent substrate (Thermo Fisher) were used for detection.

## Results

### A heat-soluble fraction isolated from *E. coli* inhibits tau aggregation *in vitro*

A common method for purifying expressed recombinant tau from *E. coli* cells takes advantage of the high solubility of tau relative to other proteins during heat treatment (13). The cell lysate is boiled to precipitate the majority of denatured proteins which are then pelleted by centrifugation - soluble tau partitions into the supernatant fraction for additional purification by cation exchange and/or size exclusion chromatography (SEC) procedures (38) (**Fig. 1A**). We attempted to optimize a rapid tau protein purification process compatible with *in vitro* aggregation assays, similar to protocols reported by Krishnakumar and Gupta (39). They observed that direct boiling of *E. coli* cells followed by centrifugation was sufficient to isolate tau protein compatible with downstream aggregation assays. Their tau fraction isolated immediately after a boiling step generated similar amyloid aggregation kinetics as monitored by Thioflavin T (ThT) when compared to tau samples further purified by affinity, ion exchange, or SEC. In our comparable protocol, proteins were expressed in Terrific Broth then sequentially processed by freeze-thaw lysis, boiling, and centrifugation to isolate soluble tau in the clarified supernatant (which we term the heat-soluble fraction). However, when the heat-soluble fraction was used directly in a standard aggregation assay with heparin as an accelerant it produced high background ThT fluorescence values (400 RFU ± 100) with no further increases in fluorescence observed over a period of 24 hrs (**Fig. 1B**). In comparison, when the heat-soluble fraction was further purified by cation exchange chromatography and dialyzed to remove excess salt, the resulting purified tau produced characteristic aggregation kinetic curves with minimal background fluorescence readings (3 RFU ± 0.5) in the absence of heparin and a final amplitude change of 300 RFU ± 50 over the course of the reaction (**Fig. 1C**). We hypothesized that the heat-soluble fraction contains an inhibitor of tau aggregation that is heat resistant, does not bind the cation exchange resin and could be isolated in the flow-through (FT) fraction. We confirmed that the presence of the FT fraction dose dependently inhibited the aggregation of 10 µM purified tau (**Fig. 1D, Supp. Fig.1A**). At concentrations of 1 % (v/v) or greater, the FT fraction did not generate ThT values that differed significantly from baseline readings. We next repeated the isolation of the FT fraction from BL21(DE3) cells transformed with an empty pUC19 vector. The FT fraction isolated from pUC19 transformed cells behaved similarly to cells transformed with pET28 0N4R tau - the ThT fluorescence did not increase over the course of the reaction in the presence of heparin, suggesting tau aggregation was inhibited (**Fig. 1E, Supp. Fig. 1B**).

**Figure 1.**
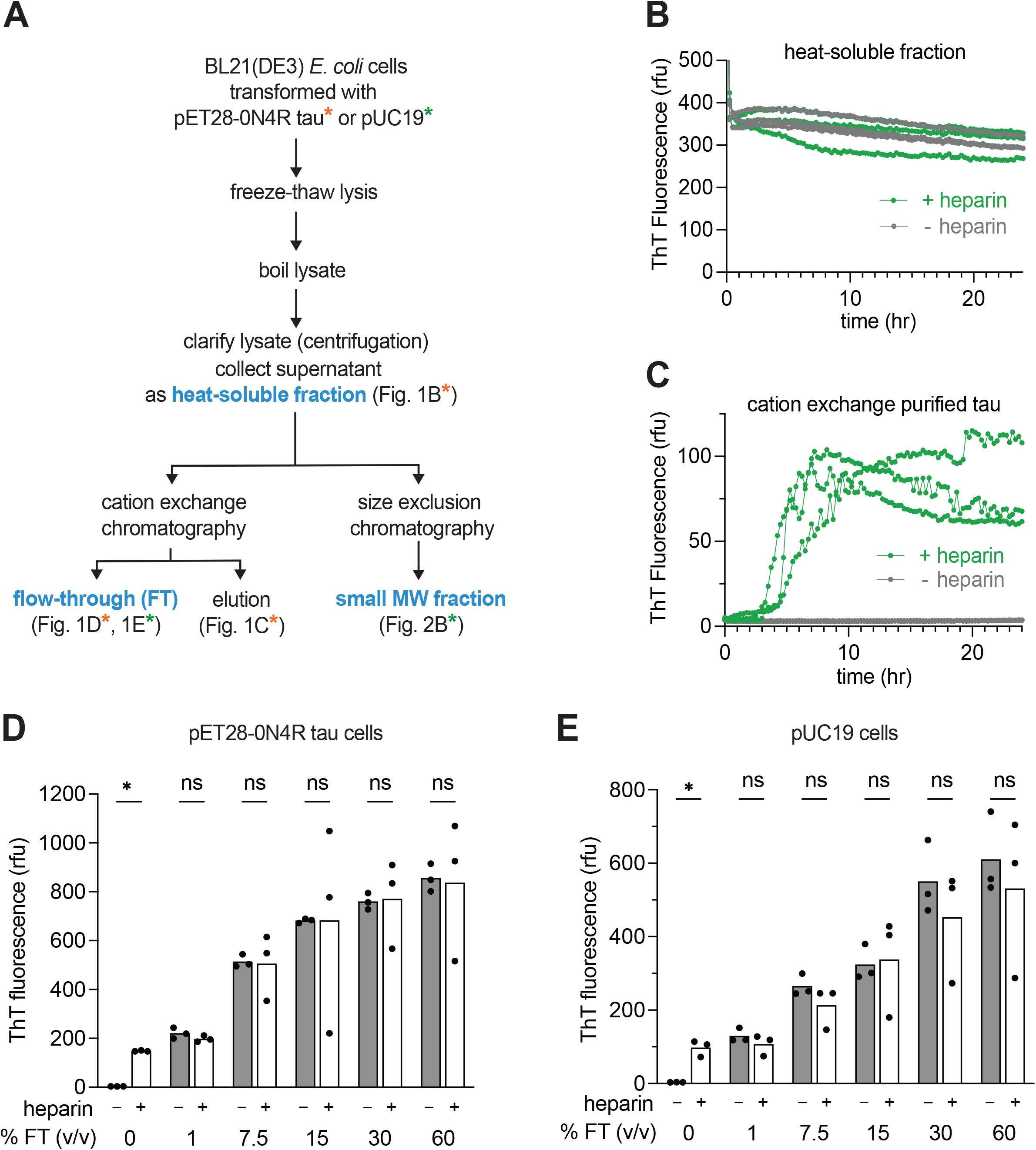
Evidence of a tau aggregation inhibitor present in *E. coli*. A) General schematic of *E. coli* fractionation procedures used in this study. Presence of inhibitor activity in tested fractions are highlighted in blue and corresponding results are shown in the figure number indicated. Colored asterisks beside figure numbers indicate if the fraction used was derived from BL21 (DE3) cells transformed with puC19 or pET28 0N4R tau vector. B-C) Aggregation kinetics of tau (10 µM) isolated in the B) heat-soluble fraction or C) after further cation exchange chromatography and buffer exchange (standard purification). Tau aggregation was monitored by ThT fluorescence signal after the addition of the accelerant heparin (green) or assay buffer (grey). For each group mean ± SD is plotted (n=3 replicates), representative of 3 independent experiments. D-E) ThT fluorescence values after 24 hrs aggregation reaction with purified tau (10 µM) in the presence of the FT fraction (%, v/v), in the absence (-) or presence (+) of heparin. D) Isolate from cells transformed with pET28 0N4R tau or E) pUC19. Individual values (n=3) and mean (bars) plotted, p<0.05, unpaired t-test, for multiple comparisons, the Holm-Šídák method was used to adjust p-values, representative of 3 independent experiments.

To rule out that the high ThT background inherent to the FT fraction was not masking amyloid detection in our assays, aggregation reactions carried out in the presence of heat soluble pUC19 lysate were processed by high-speed centrifugation and the resulting supernatant and pellet fractions were analyzed by immunoblot for total tau. The presence of heat-soluble pUC19 lysate during aggregation reduced the total amount of pelletable tau confirming decreased tau aggregation **(Supp. Fig. 1B)**. The C-terminal tau antibody (D1M9X) used to probe for tau detected full-length tau in both the pellet and supernatant fractions with minimal degradation products observed in either fraction supporting that protease digestion of tau could not account for loss of its aggregation.

### Isolation and identification of methylphosphonic acid as a tau aggregation inhibitor

To further purify and characterize the aggregation inhibitor, we separated the BL21(DE3)-pUC19 derived heat-soluble fraction by SEC which generated the 280 nm elution profile in Fig. 2A. Peak fractions (1 mL each) collected from the void volume (40 mL) to 1.4 column volumes (170 mL) were assayed for tau anti-aggregation activity. The baseline subtracted amplitudes for ThT fluorescence are reported in Supp. Table 1. Fractions at retention volumes from 103 to 108 mL significantly decreased the ThT fluorescence values by 50 ± 5% when compared to controls (**Fig. 2B, C**). The 103 and 108 mL fractions also generated minimal background fluorescence on their own suggesting effective separation of interfering signal components (Supp. Table 1). The fractions with a retention volume of 103-108 mL corresponded to a MW of less than < 2.5kDa, near the end of the total column volume (120 mL). We further tested fractions with retention volumes surrounding 103 mL in our aggregation assay and pooled all fractions with inhibitory activity which we termed the “small MW fraction”. We probed for protease-resistance of tau aggregates in completed aggregations reactions (24 hrs). Tau amyloid aggregates contain a regularly packed core region that is resistant to protease digestion which gives rise to characteristic protein fragments that can be separated by gel electrophoresis (36). Trypsin digestion of our WT 0N4R tau aggregates generates protease-resistant fragments with major peak intensities calculated at 10, 16, and 24 kDa. The presence of the SEC fraction in tau aggregation reactions decreased the peak heights corresponding to all three protease-resistant fragments (**Fig. 2D**).

**Figure 2.**
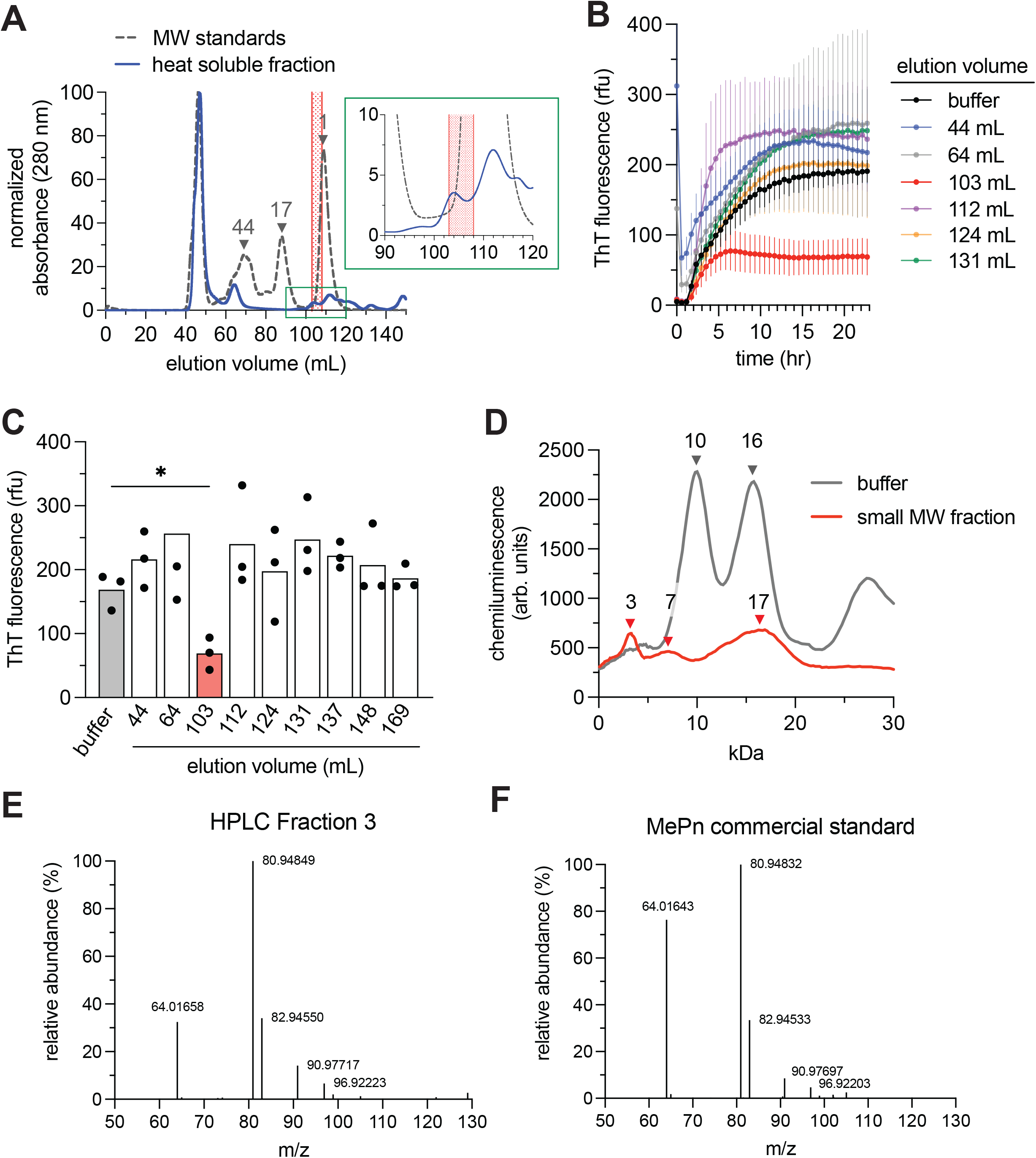
Fractionation of *E. coli* lysate to isoalte the minimal component(s) with inhibitory activity. A) Chromatogram of heat-soluble fraction from pUC19 transformed BL21 (DE3) cells subject to size exclusion chromatography. The absorbance at 280 nm of eluted fractions is plotted (solid line). The standards (kDa) indicated by arrows. Fractions from the region highlighted in red showed inhibitory activity in the tau aggregation assay. The area highlighted by the green box is enlarged in the inset graph to show the elution profile in the region with inhibitory activity. B) Aggregation kinetics of purified tau (10 µM) in the presence of 60% (v/v) of fractions collected at the indicated elution volumes. Reaction progress monitored by ThT fluorescence plotted as mean ± SD (n=3), representative of 3 independent experiments. C) Plot of endpoint ThT fluorescence values from reactions shown in B). Individual values (n=3) and mean (bars) plotted, *p < 0.05, one-way ANOVA, post-hoc Dunnett’s T3 test. D) Capillary gel electrophoresis of trypsin digested tau aggregation reactions. Chromatogram for total protein detection of trypsin-resistant fragments profiles by MW (kDa). The MW of major fragment peaks are indicated by arrows. Results representative of 3 independent experiments. E-F) Direct-infusion ESI-Q Exactive Orbitrap MS for C) HPLC fraction 3 and D) commercially available MePn (98%).

The small MW fraction was directly analyzed by one dimensional (1D) proton NMR with the resulting NMR spectrum displayed in Supp. Fig. 2A. A prominent doublet peak was detected in the spectrum with a chemical shift centered at 1.16 ppm, as well as several less intense peaks between 3.5-4.1 ppm and between 0.8-1.5 ppm. Focusing on the doublet centered at 1.16 ppm, we initially looked for a proton causing the splitting in the 1H NMR spectrum showing the expected 1:3:3:1 quartet pattern. Not finding one, we wondered what nucleus might cause the splitting and thought about 31P as a possibility. This led to the hypothesis that this peak potentially came from MePn. We used high performance liquid chromatography to further separate the small MW fraction and analyzed the obtained fractions (fractions 1-3) by NMR (**Supp. Fig. 2B, C**). The isolated fraction (fraction 3) with the prominent doublet peak (1.0-1.5 ppm) retained inhibitory activity. In addition, the peaks between 0.8-1.5 ppm contained in an inactive fraction (fraction 1) could be attributed to methyl protons of amino acids like alanine, leucine, valine and isoleucine (**Supp. Fig. 2C**). When we analyzed the active fraction (fraction 3) by mass spectrometry it gave rise to a strong peak at 81 m/z (80.948) (**Fig. 2E**). Commercially available MePn (98%) also generated a peak at 81 m/z (80.948) (**Fig. 2F**). These results were unexpected given the calculated theoretical peak of MePn is 96 m/z. However, a major peak at 81 m/z is also observed by mass spectrometry for MePn obtained from a public standard reference database (40). The discrepancy in the mass detected is likely due to loss of the methyl group when subject to high voltage ionization. Given the biophysical similarities between our sample and commercial MePn we proceeded with testing the anti-aggregation activity of MePn in our tau aggregation assays.

MePn typically has two pKa values: one at 2.1 and another one at 7.3 (41). Dissociation of both the protons are required to achieve the buffering range at around pH 7.4 where we perform the kinetic assays to mimic physiological pH conditions. When the MePn stock solution was adjusted to pH 7.4, then added to tau aggregation assays, it caused a dose-dependent reduction in the final ThT fluorescence values at 24 hrs (**Fig. 3A**) with 260 mM MePn reducing the signal by 91 ± 6%. MePn also reduced the total amounts of pelletable tau aggregate material (**Fig. 3B**) and protease-resistant tau (**Fig. 3C**) recovered from aggregation assays.

**Figure 3.**
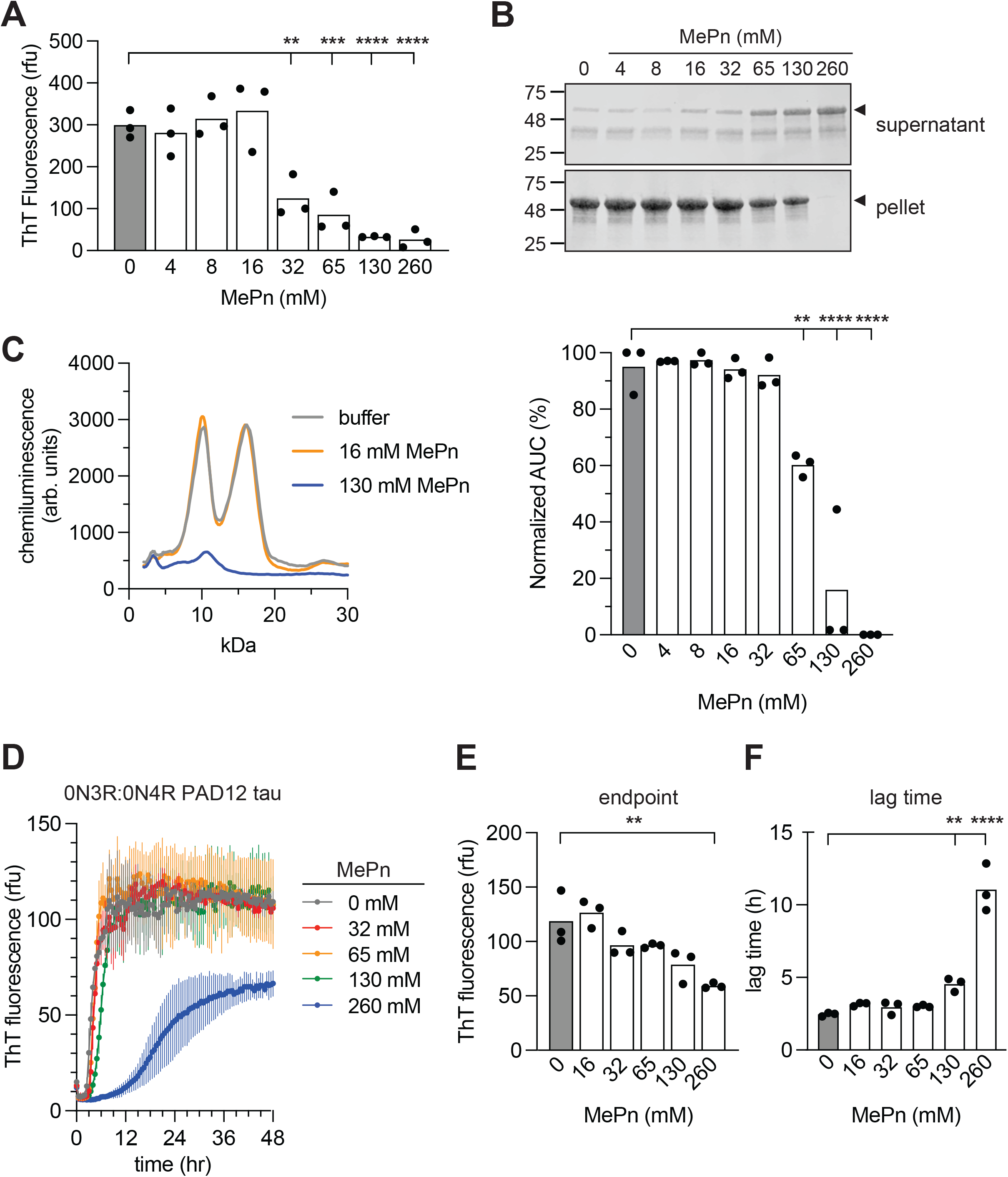
Methylphosphonic acid (MePn) inhibits *in vitro* tau aggregation. A) ThT fluorescence values after 24 hrs aggregation reaction with purified tau (10 µM) in the presence of MePn at the concentrations indicated. Mean ± SD is plotted (n=3), **=p<0.01, ***=p<0.001, ****=p<0.0001, Dunnett’s test used, representative of 3 independent experiments. B) Top: Coomassie-stained SDS-PAGE gel image of supernatant and pellet fractions from reactions of aggregated tau following separation by 100,000 x g centrifugation. Aggregation reactions were carried out with MePn at the concentrations indicated. Bottom: Quantification of densitometric signals for pelleted tau plotted as mean ± SD for 3 independent experiments.**=p<0.01, ***=p<0.001, ****=p<0.0001, one-way ANOVA, post-hoc Dunnett’s test. C) Capillary gel electrophoresis of trypsin digested tau aggregation reactions in the presence of MePn. Chromatogram for total protein detection of trypsin-resistant fragments profiles by MWW (kDa). Results representative of 3 independent experiments. D) Equimolar mixtures of 0N3R PAD12 and 0N4R PAD12 tau (40 µM total) were aggregated in the presence of MePn at the concentrations indicated. Reaction progress monitored by ThT fluorescence. D) Aggregation kinetics E) final ThT fluorescence values and F) lag times for reactions are plotted as mean ± SD (n=3), representative of 3 independent experiments. *p < 0.055, one-way ANOVA, post-hoc Dunnett’s test.

Our *in vitro* tau aggregation assays are initiated by the addition of the polyanion heparin to the reaction. One possibility is that the ability of MePn to inhibit tau aggregation is limited to this specific aggregation accelerant. We set out to test if MePn could inhibit tau aggregation in the context of accelerant-free reaction conditions. The aggregation of tau can be promoted through the introduction of a defined set of 12 phosphomimetic mutations (T181D, S202D, T205D, T212D, S214D, T217D, T231D, S235D, S396D, S400D, T403D, S404D) into the 0N4R or 0N3R tau sequence to generate the variants named 0N4R PAD12 and 0N3R PAD12, respectively (35, 42). Aggregation of an equimolar mixture of 0N3R and 0N4R PAD12 (40 µM total), at 37°C, with shaking, triggered amyloid formation in the absence of any inducer (**Fig. 3D**) as reported previously (35). Under these conditions, MePn inhibited the final ThT fluorescence values after 24 hrs of aggregation although high concentrations (260 mM) were required for significant inhibitory activity (**Fig. 3E**). In contrast to heparin-induced 0N4R tau aggregation reactions, MePn also significantly delayed the lag time to amyloid formation for 0N3R:0N4R PAD12, at concentrations of ≥130 mM MePn (**Fig. 3F**). Similar inhibition effects were observed with MePn in aggregation reactions of 40 µM 0N4R PAD12 (**Supp. Fig. 3**).

### Culture media containing methylphosphonic acid is sufficient to inhibit tau aggregation in *E. coli*

Certain bacteria such as marine bacteria have the ability to both synthesize and degrade MePn as a source of phosphate (43, 44). *E. coli* do not possess the pathways to synthesize MePn. However, they are capable of taking up and metabolizing phosphonates including MePn as a source of phosphorus via a carbon-phosphorus lyase pathway genetically encoded by the *phn* operon (43). Since *E. coli* are not able to synthesize MePn we reasoned that the culture media contains MePn which is then taken up by the bacteria. We typically culture *E. coli* in TB which is a complex, nutrient-rich, buffered media that allows for increased culture densities during protein expression experiments. However, more strictly defined, minimal nutrient, formulations such as minimal media (M9) can also be used for recombinant protein expression in *E*.*coli*. When *E. coli* expressing puc19 plasmid were cultured in TB versus M9 media prior to lysis and isolation of the heat-soluble fraction as in **Fig. 1D**, only isolates from *E. coli* grown in TB media were able to significantly inhibit tau amyloid formation (48± 5%) when supplied at a final concentration of 30% (v/v) to reactions respectively (**Fig. 4A**). When supplemented at 60% (v/v) both M9 and TB grown lysates inhibited tau aggregation (**Supp. Fig. 4A**). Our results raised the possibility that MePn in the culture media was being internalized by *E. coli* to generate an environment that inhibits tau aggregation. Thus, we assessed the ability of culture media-derived MePn to prevent tau aggregation in living *E. coli* cultures. Previous reports demonstrated that when *E. coli* cultures were grown in M9 media and induced to overexpress the human tau protein, tau accumulated at high concentrations sufficient to trigger its aggregation in the live bacteria (37, 45, 46). We also aimed to assess the specificity of MePn in modulating the amyloid formation propensities, whether it is restricted to tau or extends its inhibitory activity to other amyloidogenic proteins like polyQ-expanded huntingtin (mutant Htt) (47–49). *E. coli* grown in M9 media were induced to express 0N4R tau or His and MBP tagged mutant Htt construct containing the exon 1 sequence of Htt with a 46Q expansion. Tau and mutant Htt amyloid formation in *E. coli* was quantified by the Thioflavin S (ThS) fluorescence which has similar amyloid binding properties as ThT. Expression of either protein for 18 hrs led to increased fluorescence signal intensity for ThS across the emission range measured (475-600 nm) as compared to uninduced culture controls (**Fig. 4B-D**). For cells expressing 0N4R tau, supplementing MePn to the M9 media for the duration of protein expression resulted in a dose-dependent decrease in the ThS signal, that was significant with 16 mM MePn and at 130 mM MePn reduced the amyloid signal by 50 ± 10% compared to the 0 mM MePn control group. Mutant Htt amyloid formation was resistant to inhibition by MePn supplementation as 130 mM MePn was required to achieve significant inhibition (**Fig. 4C**). The amount of tau or mutant Htt protein expression was unchanged between *E. coli* cultures grown with or without the concentrations of MePn tested indicating that the MePn is preserving the high concentration of intracellular tau in a non-aggregated state (**Supp. Fig. 4B-C**). Thus, our results support that MePn in the *E. coli* culture media is also sufficient to inhibit tau aggregation in living cells (43).

**Figure 4.**
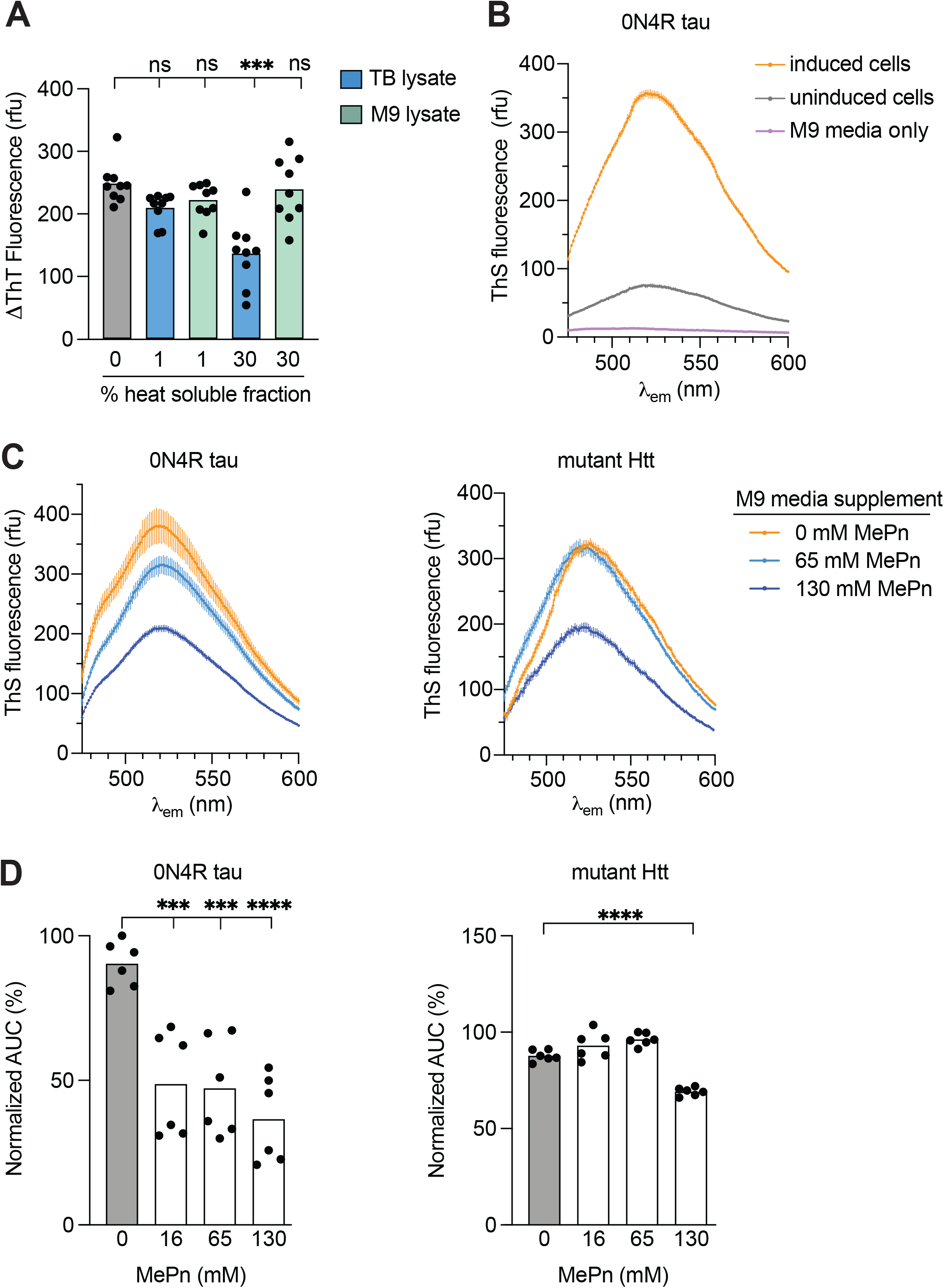
Supplementing MePn to the culture media inhibits tau aggregation in *E. coli*. A) Aggregation kinetics of 0N4R tau (10 µM) in the presence of 1%, v/v of small MW fraction isolated from TB (green bar) or M9 (blue bar) compared to a buffer only control (grey bar). Difference in ThT fluorescence values between and initial and end time points plotted as mean ± SD (n=9), ****p <= 0.0001, one-way ANOVA, post-hoc Dunnett’s T3 test, representative of 9 independent experiments. B) Thioflavin S measurement of amyloid formation in *E. coli* cells before (uninduced cells) or after (induced cells) 18 hrs of 0N4R tau protein expression. ThS fluorescence values plotted as a function of emission wavelength, mean ± SD (n=3), representative of 3 independent experiments. The fluorescence spectrum of M9 culture medium (control) is also plotted. C-D) Top: *E. coli* cells expressing C) 0N4R tau or D) mutant Htt cultured in M9 media supplemented with the indicated concentrations of MePn and processed as described in B). Bottom: Plot of normalized AUC of baseline-subtracted experimental groups. For each group mean ± SD is plotted (n=3 replicates), representative of 3 independent experiments.

## Discussion

In this study, we isolated and characterized a fraction from *E. coli* lysate that inhibits tau aggregation. The inhibitory fraction consisted of several small MW components with initial analysis leading us to test the inhibitor candidate methylphosphonic acid (MePn). We verified that MePn inhibits tau aggregation *in vitro*. Furthermore, we showed that supplementing MePn to the culture media was sufficient to inhibit tau aggregation in *E. coli, in cellulo*.

Our study has generated important insights for tau production and tau aggregation studies in *E. coli* experimental systems. We confirmed previous reports that overexpression of human tau can lead to aggregation in live *E. coli* cultures (37, 45). Importantly, we uncovered that the degree of tau aggregation occurring in *E. coli* can be readily controlled by modifying the culturing media. M9 media, which is strictly defined and is relatively nutrient-poor compared to other media used for *E. coli* cultures (e.g.

Luria Broth and TB). Our results support that M9 media generates a cellular environment which allows for tau monomers expressed at high concentration to self-associate and aggregate on a short time scale (<18 h). Therefore, we recommend culturing in M9 media for *in vivo* screening platforms to identify modulators of tau aggregation. However, due to the significant generation of aggregates, culturing *E. coli* in M9 media for the purposes of purifying tau protein monomers is not advisable unless an effective denaturing step such as boiling or detergent is included in the purification procedure. In contrast, our results with *E. coli* cultured in TB suggest that this rich media formulation contains components that directly inhibit tau aggregation. Additional purification steps are needed to separate anti-aggregation components as well as background fluorescent factors from tau protein extracts and allow for downstream aggregation assays. Our results on the differential effects of culture media on tau aggregation may provide an explanation of why we were unable to replicate a previous report that tau protein is competent to aggregate *in vitro* following simplified tau protein isolation procedures from *E. coli* (39). The finding that MePn supplemented to the media was able to inhibit amyloid formation by tau in *E. coli* cultures at much lower concentrations than required for inhibition of mutant Htt indicates increased specificity or sensitivity of MePn towards tau aggregation. It is possible that other yet identified metabolites may be more efficacious against mutant Htt aggregation in cells.

A limitation of our study is that we were not able to fully characterize the composition of our small MW fraction. Our initial analysis led us to test MePn however, high concentrations of MePn were required to inhibit aggregation *in vitro* leading us to suspect that as yet, unidentified components in the fraction are also contributing to the inhibitory activity we observe. Our identification of amino acids in inactive fractions by NMR could indicate the presence of a peptide carrier for MePn that might facilitate aggregation inhibition and account for the unexpected SEC retention volume correlating with larger molecules (2.5 kDa). Another limitation of our discovery that MePn inhibits tau aggregation in *E*.*coli* cultures is that the effect may be mediated by both direct and indirect downstream actions. For example, it is possible that metabolism of MePn by the carbon-phosphorus lyase pathway (43) could generate intermediates that also inhibit tau aggregation. Further investigations manipulating relevant metabolic pathway genes or refining the fractionation of candidate aggregation inhibitors will contribute to resolving this outstanding question.

Human tau contains three characterized aggregation motifs (50, 51) but remains highly soluble due to the abundance of charged residues distributed throughout the protein sequence that limits self-association. Polyanions share characteristic properties to 1) interact with positively charged tau residues and 2) scaffold multiple tau molecules in close proximity to promote aggregation (31, 52). With respect to inhibiting tau aggregation in cells, many of the identified factors are protein partners such as molecular chaperones (53) or polymerized microtubule proteins (54) that in effect limit tau-tau interactions. PTMs on tau have also been shown to inhibit (29) or promote (52–54) its aggregation depending on the specific modifications introduced. Currently, we find only one example in the literature of a native cellular metabolite or small biomolecule in human cells that can directly inhibit tau aggregation - phosphatidylserine has been shown to inhibit tau aggregation *in vitro* with its activity being dependent on the length and degree of saturation of the attached fatty acids chains (55). Biosynthetic pathways for MePn and other phosphonates are generally restricted to microorganisms that utilize phosphonates as a source of phosphorus (56, 57). However, MePn can also be found in human cells in the very limited circumstance of exposure to nerve agents where MePn is formed as a degradation product (57). MePn is not cytotoxic to cells when tested at low micromolar concentrations (58). Future studies investigating metabolites have the potential to reveal a pool of factors with similar properties to MePn that function in healthy cells to maintain tau in a soluble state.

## Acknowledgements

We thank the Alberta Proteomics and Mass Spectrometry Facility (University of Alberta), NANUC (University of Alberta), and the Dept. of Chemistry (University of Alberta) for access to instrumentation used in this study. We thank R. Fahlman for critical discussions of the project during its development. This work was supported by NSERC Discovery Grant funding to S.A.M (RGPIN-2019-06230).

## Author Contributions

M.S., A.Y., and S.A.M. conceived and designed the study. M.S., A.P., T.P., J.M., E.G., A.Y., B.S., O.J., and S.A.M. acquired, analyzed, or interpreted data. M.S. and S.A.M. drafted and revised the manuscript.

## Conflict of Interest

The authors declare that they have no conflicts of interest with the contents of this article.

## Footnotes

### Abbreviations

DTT: Dithiothreitol
E. coli: Escherichia coli
FT: Flow-Through
G6P: Glucose 6-phosphate
HPLC: High Performance Liquid Chromatography
IPTG: isopropyl-beta-D-thiogalactopyranoside
MAPT: microtubule associated protein tau
MePn: Methylphosphonic acid
MS: Mass Spectrometry
NMR: Nuclear Magnetic Resonance
PAD12: 12 phosphomimetics of AD
PMSF: phenylmethylsulfonylfluoride
PTMs: Post-Translational Modifications
R5P: Ribose 5-Phosphate
SEC: Size Exclusion Chromatography
TB: Terrific Broth
ThS: Thioflavin S
ThT: Thioflavin T

## Notes

### Competing Interest Statement

The authors have declared no competing interest.

