## Supplemental figures and table for "A metabolite extracted from *E. coli* suppresses tau aggregation"

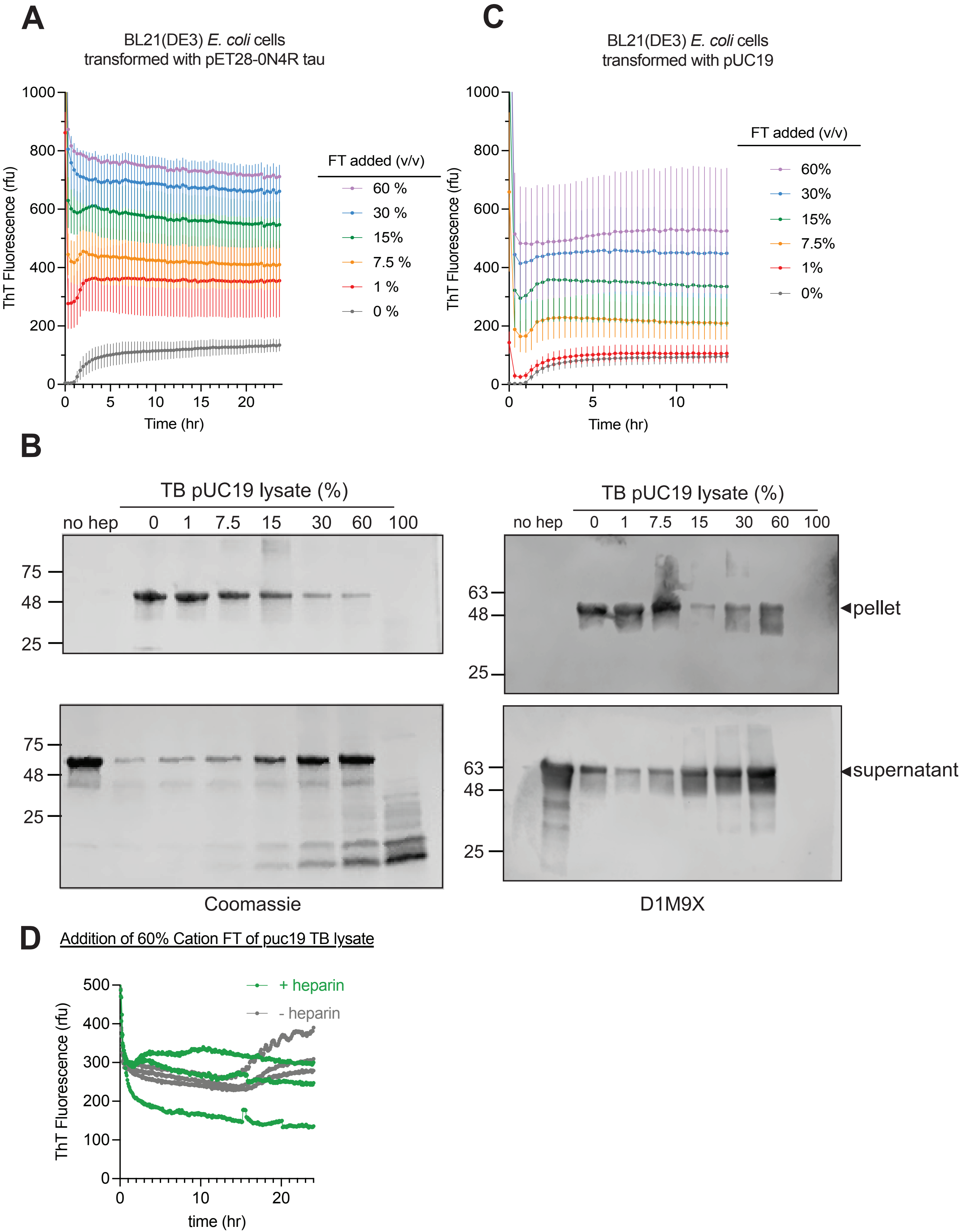

A
BL21(DE3) E. coli cells
transformed with pET28-0N4R tau
ThT Fluorescence (rfu)
Time (hr)
FT added (v/v)
60 %
30 %
15%
7.5 %
1 %
0 %
C
BL21(DE3) E. coli cells
transformed with pUC19
ThT Fluorescence (rfu)
Time (hr)
FT added (v/v)
60%
30%
15%
7.5%
1%
0%
B
TB pUC19 lysate (%)
no hep
0
1
7.5
15
30
60
100
75
48
25
63
◄pellet
◄supernatant
Coomassie
D1M9X
D
Addition of 60% Cation FT of puc19 TB lysate
+ heparin
- heparin
ThT Fluorescence (rfu)
time (hr)

**Supplementary Figure 1:** Supporting data for Figure 1. Aggregation kinetics of tau (10 µM) in the presence of FT fraction (%, v/v) isolated from cells transformed with A) pET28 0N4R tau or C) pUC19. Tau aggregation was monitored by ThT fluorescence signal after the addition of the accelerant heparin. For each group mean ± SD is plotted (n=3 replicates), representative of 3 independent experiments. B) Coomassie-stained SDS-PAGE gel image and D1M9X immunoblot of supernatant and pellet fractions from reactions of aggregated tau following separation by 100,000 x g centrifugation. Aggregation reactions were carried out with TB pUC19 lysate at the concentrations indicated. D) Aggregation kinetics of 0N4R tau (10 µM) in presence of 60% TB pUC19 lysate. Reaction progress monitored by ThT fluorescence plotted as mean ± SD (n=3), representative of 3 independent experiments.

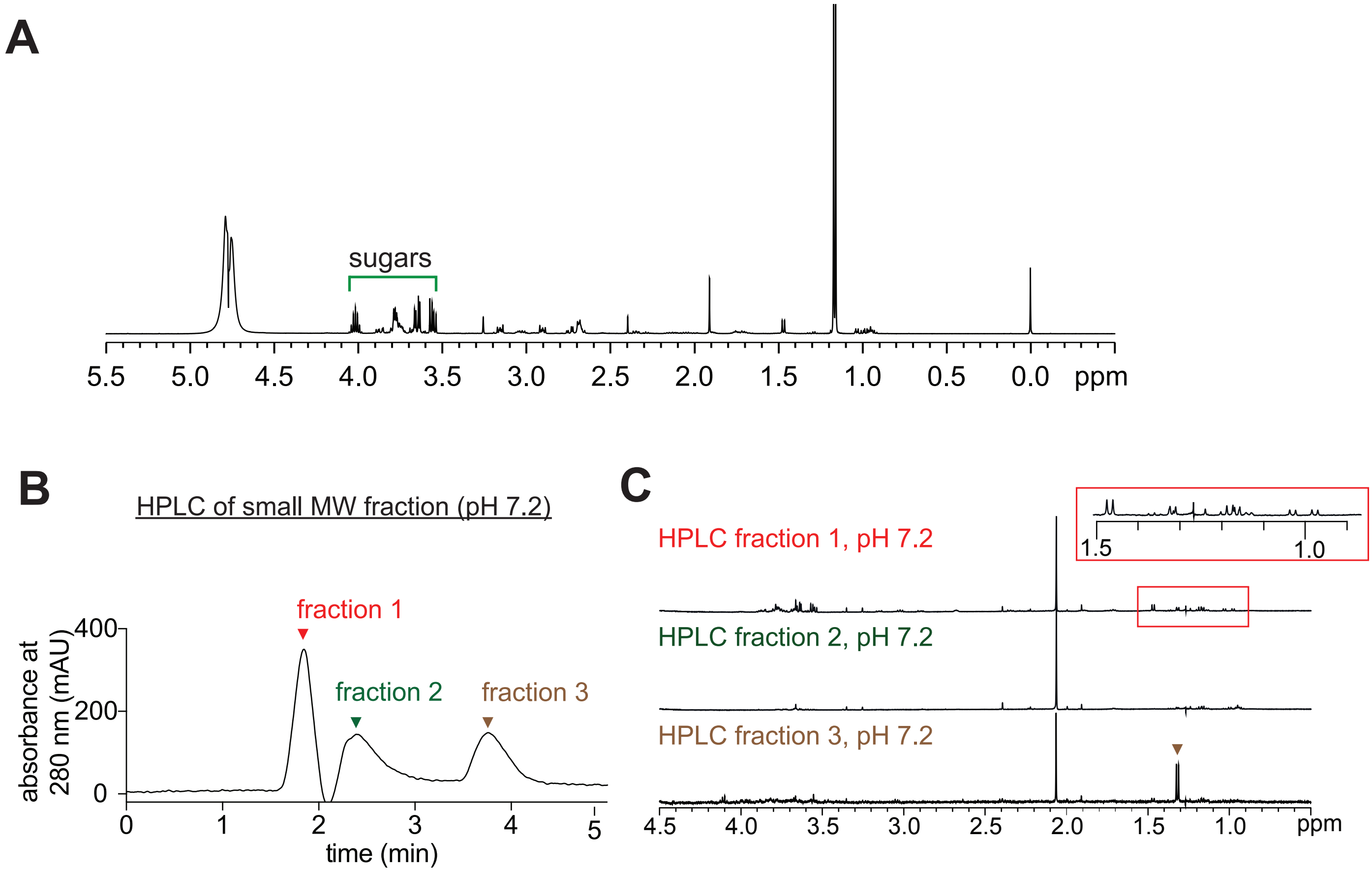


**Supplementary Figure 2:** Supporting data for Figure 2. A) 1D 1H NMR spectrum of the small MW fraction in D2O (pH 7.2) acquired at 500 MHz. Major peaks are indicated by arrows. Green arrow indicates peaks potentially matching with MePn. Red arrows indicate peaks corresponding to amino acids. B) Absorbance profile at 280 nm of small MW fraction separated by HPLC. The elution corresponding to the three major peaks indicated were collected as fractions 1-3 (labeled) for further analysis. C) 1D 1H NMR spectrum of the HPLC fractions 1-3 from A) at pH 7.2. The area highlighted by the red box is enlarged in the inset graph to show the amino acid peaks. Peaks predicted to correspond to MePn (brown) are indicated by the arrowheads.

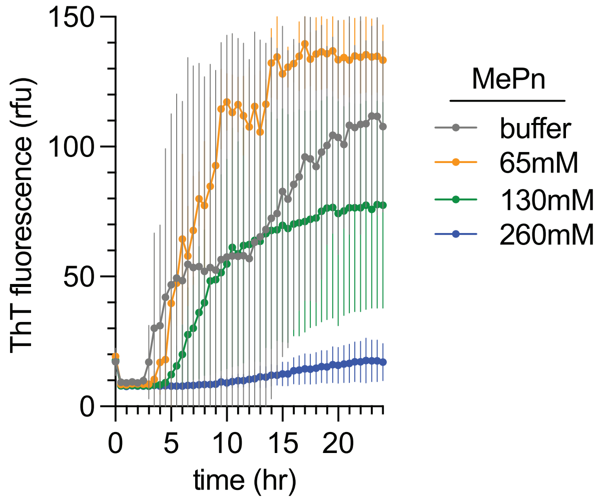


**Supplementary Figure 3:** Supporting data for Figure 3. Aggregation kinetics of 0N4R PAD12 (40 μM) in the presence of the indicated concentrations of methylphosphonic acid. Reaction progress monitored by ThT fluorescence plotted as mean ± SD (n=3), representative of 3 independent experiments.

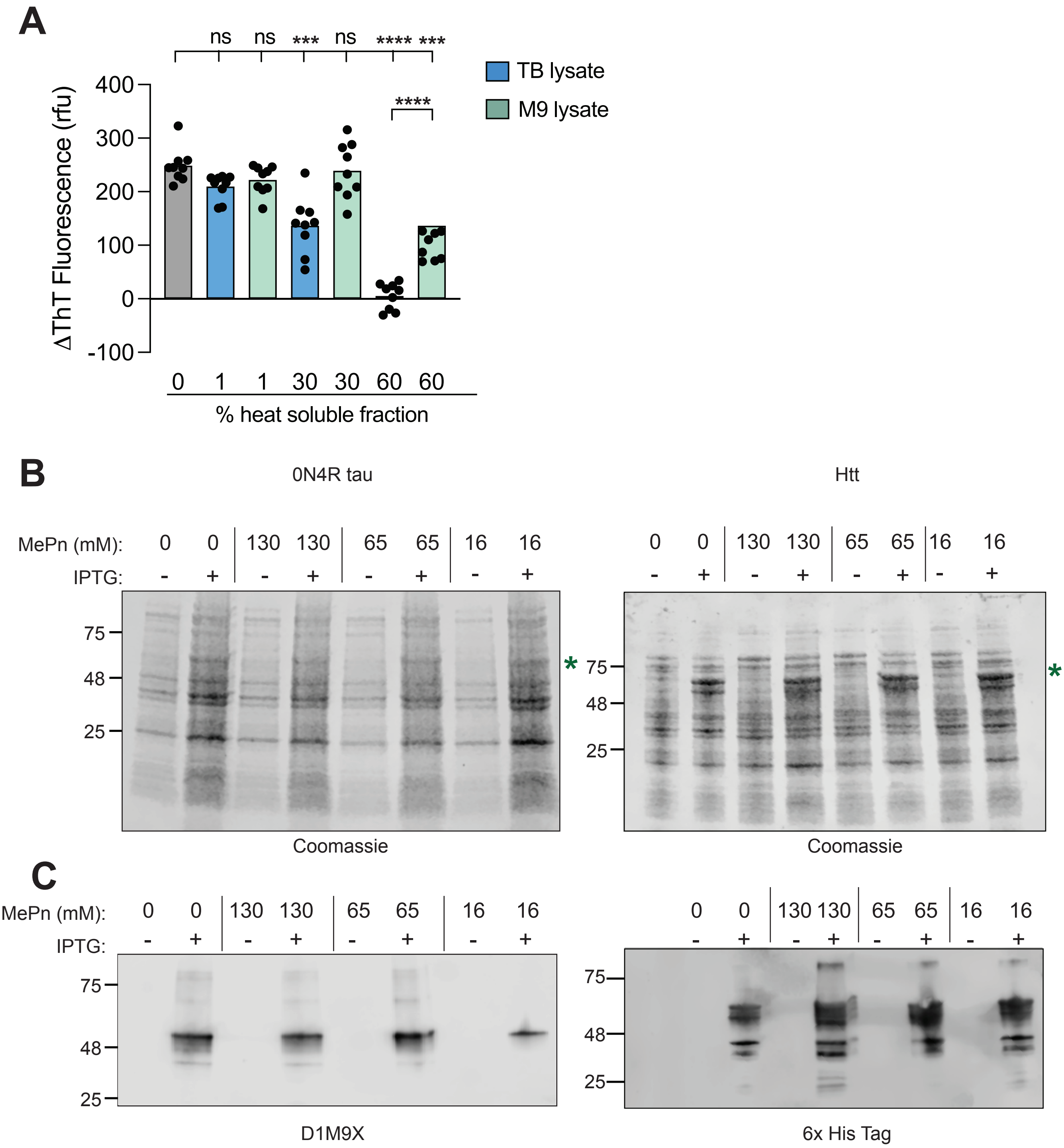
A
ns
ns
***
ns
****
***
****
TB lysate
M9 lysate
ΔThT Fluorescence (rfu)
400
300
200
100
0
-100
0 1 1 30 30 60 60
% heat soluble fraction
B
0N4R tau
Htt
MePn (mM): 0 0 130 130 65 65 16 16
IPTG: - + - + - + - +
75
48
25
*
Coomassie
C
MePn (mM): 0 0 130 130 65 65 16 16
IPTG: - + - + - + - +
D1M9X
6x His Tag

**Supplementary Figure 4:** Supporting data for Figure 4: A) Aggregation kinetics of purified tau (10 µM) in the presence of 30 and 60% (v/v) of pUC19 lysate isolated from TB (green bar) or M9 (blue bar) compared to a buffer only control (grey bar). Difference in ThT fluorescence values between and initial and end time points plotted as mean ± SD (n=9), ****p <= 0.0001, one-way ANOVA, post-hoc Dunnett's T3 test, representative of 9 independent experiments. B) SDS-PAGE of *E. coli* lysates expressed 0N4R tau (left) or mutant Htt (right) cultured in M9 media in the presence (+) of indicated concentrations of MePn or buffer only (-) as controls. Tau and mutant Htt protein expression was induced by the addition of IPTG (+) and compared to samples grown without IPTG (-). Green asterisk (*) indicates the predicted migration of tau and mutant Htt protein (His-MBP- tagged construct) observed in samples induced with IPTG. Immunoblotting with the respective antibodies revealed comparable levels of protein expression.

| **Peak Number** | **Elution Volume (mL)** | **Endpoint ThT after baseline subtraction (RFU)** |
|---|---|---|
| Buffer alone | | 191.886 ± 14.673 |
| 1 | 44.42 | 216.305 ± 44.155 |
| 2 | 64.42 | 256.491 ± 136.633 |
| **3** | **103.42** | **69.080 ± 25.240** |
| 4 | 112.42 | 240.155 ± 80.029 |
| 5 | 124.42 | 197.615 ± 72.812 |
| 6 | 131.42 | 247.377 ± 59.425 |
| 7 | 137.42 | 221.827 ± 20.400 |
| 8 | 148.42 | 207.158 ± 56.476 |
| 9 | 169.42 | 186.436 ± 19.985 |

**Supplementary Table 1**: Aggregation assay results for inhibitor activity of lysate fractions separated by SEC. *E.coli* cells transformed with pUC19 vector were lysed and subject to SEC. Fractions corresponding to peaks of absorbance values at 280 nm and indicated elution volumes (mL) were tested in tau aggregation assays containing 10 µM tau + 60% fraction (v/v), monitored via ThT fluorescence (RFU).
